# Similar won’t make it: Coexistence, habitat availability and suboptimal restoration

**DOI:** 10.1101/2024.07.24.604984

**Authors:** Felipe Maresca Urioste, Ana Inés Borthagaray, David Cunillera-Montcusí, Álvaro Soutullo, Matias Arim

**Author notes:** **Corresponding author:** Felipe Maresca Urioste, Tacuarembó S/N between Bvar. Artigas y Av. Aparicio Saravia, Maldonado, Uruguay. **Funding Statement:** This study was Funded by Horizon 2020 Framework Programme (Grant Number 869296) and by grants CSIC-grupos (ID 657725) and ANII FCE_1_2023_1_176573 to MA.

## Abstract

In the context of the looming biodiversity crisis, adding patches of habitat has been proposed as a restoration strategy in several ecosystems. However, these interventions rarely mimic the conditions of the natural habitats precisely. Consequently, not all species equally benefit from the restored habitat, fostering a difference in species fitness that could generate an imbalance in the equalizing and stabilizing mechanisms that enable species coexistence, leading to competitive exclusion. Using a metapopulation model, we explore how the introduction of habitat patches with biased fitness advantages affects species coexistence. We show that while adding habitat patches generally increases species occupancy, suboptimal patches can favor one species, disrupting the stabilizing and equalizing mechanisms that enable coexistence, leading to competitive exclusion. This impact on the coexistence regime is particularly pronounced in species with high niche overlap and high colonization-extinction ratios. Our results highlight the double-edged sword effect of suboptimal restoration, revealing potential unintended consequences that could exacerbate biodiversity loss. On the other hand, restoration favoring endangered species is also identified as a conservation tool. The present study advances in the mechanistic comprehension of habitat restoration, a necessary endeavor in the face of the challenges posed by global change.

## 1. Introduction

Habitat loss is considered one of the main drivers of the current biodiversity crisis (Millenium Ecosystem Assessment 2005; FAO 2019, Tickner et al., 2020; Albert et al., 2020), being identified as a cause for biodiversity loss at several organizational levels from the individual to the community (Nee & May 1992; Hanski 1994; Moilanen & Hanski 1995; Hanski 1997; Gyllenberg & Hanski 1997; Bascompte & Solé, 1998; Klausmeier 2001; Battisti et al., 2003). Habitat restoration has been proposed and encouraged in recent decades as a viable way of mitigating the problems caused by habitat loss and fragmentation by means of nature-based solutions to global change (Dobson 1997; Abrams 2018; United Nations Decade on Ecosystem Restoration Resolution, 2019; Cuenca-Cambronero et al. 2023). However, a gap between restoration actions and the comprehension of the mechanisms responsible for community assembly at the landscape scale persists (Laurance et al., 2012; Durant et al., 2019; Cooke et al., 2020; Kadykalo et al., 2021; Aoyama et al., 2022). Most papers published on the subject refer to intervention proposals and rarely to their performance, and even then, there is likely a bias towards reporting interventions with positive results (Godet & Devictor, 2018). Indeed, counterintuitive results that challenge our understanding of the biodiversity-landscape interactions have been reported (Moor et al. 2022). While a general positive effect of the addition of patches is observed, unexpected effects are also reported without a clear mechanism for explaining them (Ruhi et al., 2013; Aoyama et al. 2022; Moor et al. 2022).

Overall, meta analyses have estimated the effectiveness of this kind of measures at 80% at most, which is indeed high, but still means that they do not work as intended in at least 20% of the cases (Sutherland et al., 2015). In this context, the potential of theoretical ecology as means for advancing in mechanistic based ecological interventions has been highlighted (Perring et al., 2015; Török & Helm, 2017; Gawecka & Bascompte 2021). In a similar manner, restoration and conservation ecology provides an ideal, yet also underexploited scenario to test the predictions of ecological theory, often being referred to as their “acid test” (Young et al., 2005; Török & Helm 2017).

Modern Coexistence Theory (MCT) provides a phenomenological, high level theoretical framework that addresses the balance between mechanisms that allow the coexistence of species (Barabas, 2018; Chesson, 2018,2020; Aoyama et al. 2022). These mechanisms can be of two classes: equalizing mechanisms, which reduce fitness differences between species, and stabilizing mechanisms, which reduce the niche overlap between them (Chesson 2000b, 2020, Barabas 2018). In the MCT framework, the species overlap () is captured by the geometric average of the inter and intraspecific species feedbacks (see Chesson 2000). On the other hand, equalizing mechanisms are captured by the ratio in species performance (i), which is determined by the inter and intra specific feedbacks experienced in a given environment by each species (see Chesson 2000b, Hubbell 2001). This is captured in a simple mathematical representation of the conditions that allow coexistence between two species is (Chesson 2020):

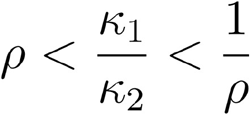

which allows for a visual representation of the parameter space where stable coexistence is possible (Figure 1).

**Figure 1.**
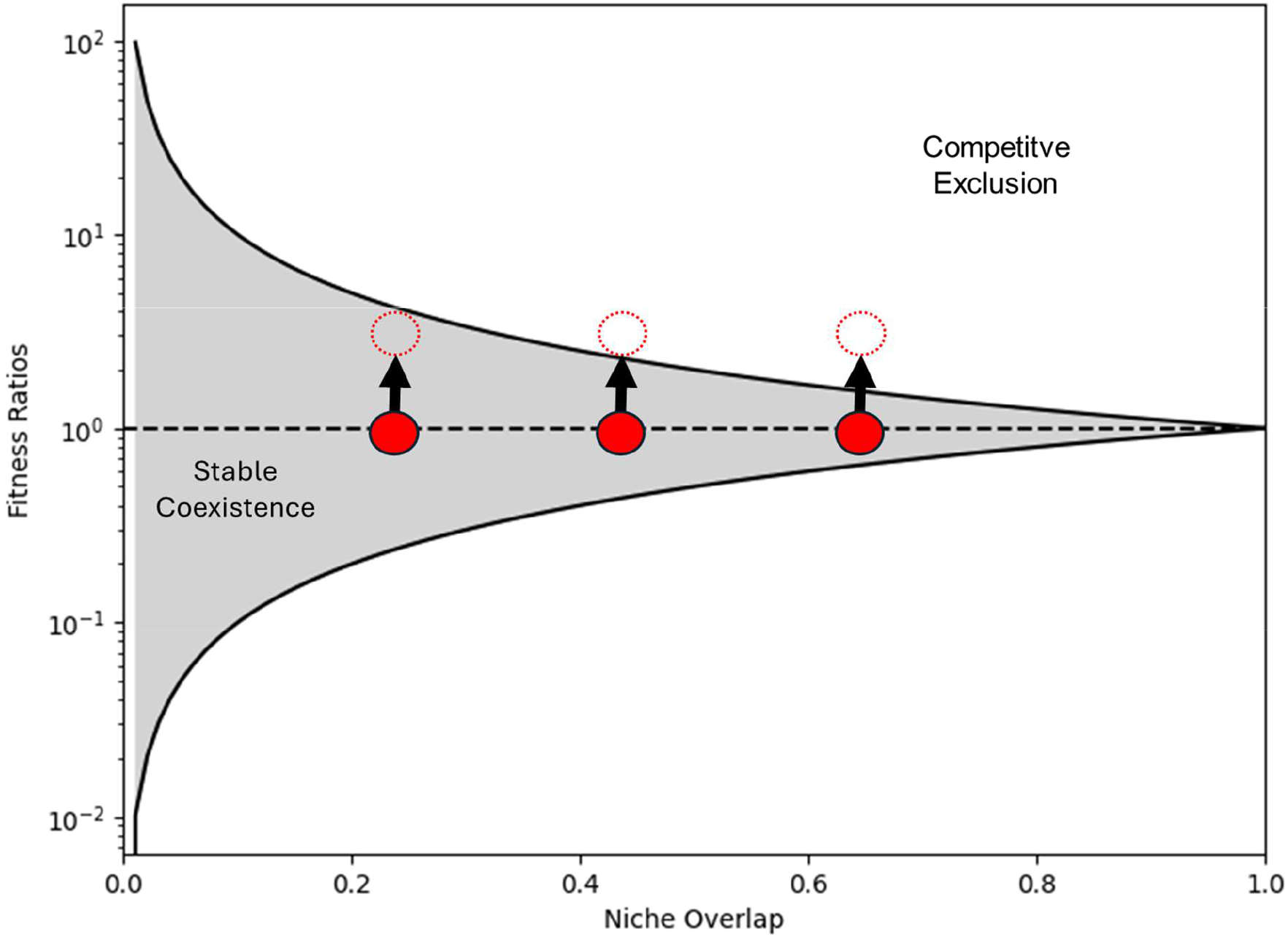
Graphical representation of the hypothesized relationship between Modern Coexistence Theory (MCT) and undesired effects of suboptimal restoration. The 2 species system analyzed in our model are represented by the red circles. Solid filled circles represent the 2 species system before the habitat is added. Adding habitat that benefits one species more than other changes the fitness ratios, thus leading to a shift in the system represented by the arrows. If this shift is suficiently large, the new location of the system could be outside the region that allows for stable coexistence, thus leading to competitive exclusion of one of the species.

Many habitat restoration measures consist in reestablishing or creating patch habitats in order to preserve viable populations (e.g. Moore et al., 2022). Adding patches of habitat prevents extinction by increasing the species metapopulation carrying capacities (Ovaskainen and Hanskii 2000, Tao et al., 2024). However, restored habitat patches may be suboptimal (Benayas et al., 2009; Crouzeilles et al., 2016; Cabrera et al., 2019; Hernandez-Carrasco et al., 2023), in the sense that they have physical or biological properties that are different from the original habitats that are being restored. This can make them more favorable for some species over others, providing better fitness to the favored species at the metacommunity level. This enhanced fitness modifies the balance between stabilizing and equalizing mechanisms that allow coexistence, which can promote the conditions for a competitive exclusion depending on the magnitude of the change in the relative performance of species in the landscape, which can lead to the competitive exclusion of the species with lower fitness in the novel habitat (Figure 1)

In this paper, we use a mean field metapopulation model to explore the impact over competing species of adding habitats. We incorporate a fitness bias in the new habitats in order to elucidate how such disturbance in the fitness ratios between species may affect the balance between coexistence mechanisms. Metapopulation models have provided great insight into the consequences of habitat loss and fragmentation and their underlying mechanisms (Levins, 1969; Nee & May 1992; Hanski 1994; Tilman et al., 1994; Gyllenberg & Hanski 1997; Bascompte & Solé, 1998; Klausmeier 2001; Melián & Bascompte 2002, Fortuna et al., 2013). Interventions of adding or removing a fraction of landscapes have been previously analyzed using metapopulation models (Levins 1969, Hanski 1994). Similarly, the effect of landscape features on species interactions has also been analyzed using metapopulation models (e.g. Roughgarden 1974; Tilman, May, Lehman, & Nowak, 1994; Chen & Hui, 2009; Borthagaray et al. 2015). Although less frequent, metapopulation models have been used to explore restoration (Gawecka & Bascompte 2021). We aim to advance the mechanistic understanding of how suboptimal restoration affects species coexistence and its implications for ecosystem conservation, an area that is still poorly understood and is of critical importance in the face of global change.

## 2. Methods

We used a set of difference equations to analyze the fraction of occupied patches by two competing species in a spatially implicit habitat. We consider a total habitat quantity equal to 1 (*H*_*total*_= 1), composed of an original habitat with a fixed quantity of ½ (*H*_*1*_=0.5), and an incorporated habitat through restoration (*H*_*2)*_, whose quantity varies. The fraction of the total habitat that is not contained by either *H*_*1*_ or *H*_*2*_ is considered inhabitable by both species. In this sense, we refer to the quantity of restored habitat *H*_*2*_ as the intervention size. The dynamics of the fraction of occupied patches by each species in each type of habitat is governed by the set of equations:

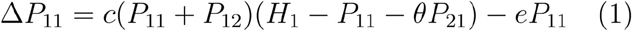

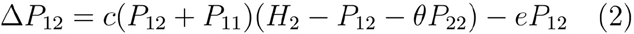

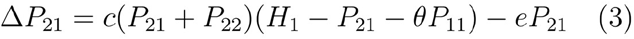

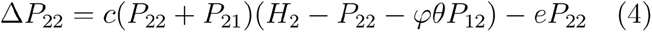

Being *P*_*ij*_ the fraction of occupied patches by species *i* in habitat *j*, and *c* and *e* colonization and extinction rates, assumed equal between competing species. *θ* is a competition parameter with values ranging from 0 to 1, with 0 being no competition and 1 being total interference. In that sense, *θ* represents the probability that a species colonizes a patch occupied by the other species. While both species interfere with each other in a symmetrical way in the original

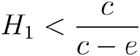

habitat *H*_*1*_, this is not necessarily the case in *H*_*2*_ (equations 3 and 4). Parameter φ, whose values also range from 0 to 1, represents the reduction in competitive interference that the species favored by the incorporated habitat (in our case species 1), has over the other in the habitat *H*_*2*_. We have called this parameter the *fidelity of the intervention*. Note that in the case where φ = 1, *H*_*1*_ and *H*_*2*_ are ecologically equivalent and the two species are equally favored by the novel environment, whereas if φ < 1, the competitive pressure that species one applies to species two is reduced in the novel environment. Equilibria for the system of equations was found by iterating numerically until all *P*_*ij*_ showed variations within a margin of 1×10^−5^. Numerical simulations were performed using values of *c* and *e* that fulfilled the condition which allows persistence in a scenario with no added habitat (Levins, 1969; Hanski 1994):

Once the equilibria was found for each set of variable parameters (*H*_*2*_, φ, *c, e* and *θ)* we proceeded to calculate the *impact of the intervention* which was defined as:

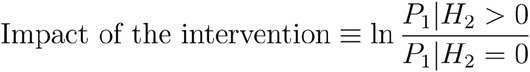

which is the log ratio between the fraction of occupied patches by the least favored species (in our case, P1) after (P1|H2>0) and before (P1|H2=0) the intervention. A negative impact of the intervention means a reduction in species occupancy. Table 1 contains a brief explanation of each of these parameters to aid in the reading.

**Table 1.**
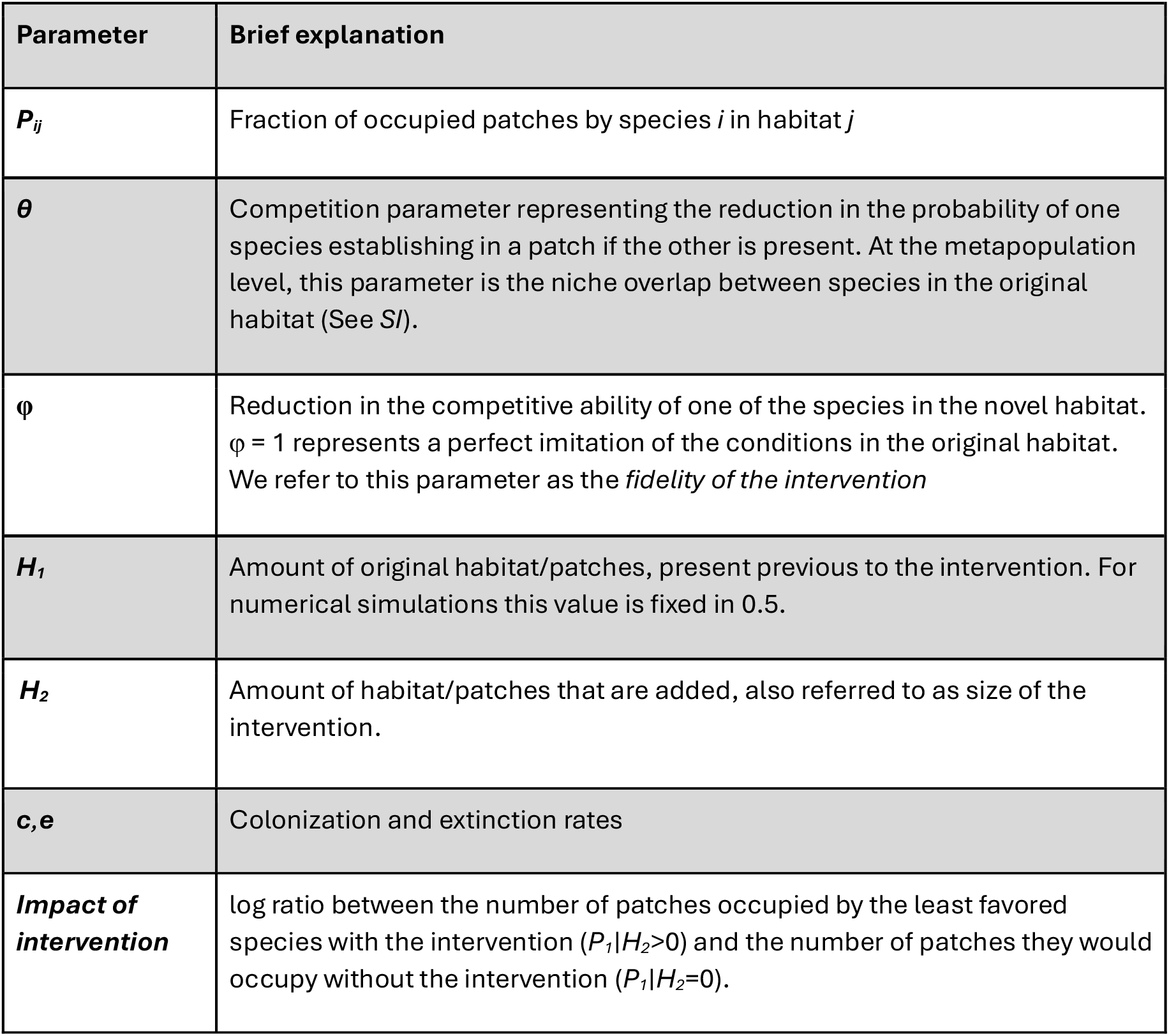
Parameters of the metapopulation model.

In order to express the results in terms compatible with the Modern Coexistence Theory, we demonstrate that parameter *θ* is equivalent to the niche overlap *ρ* prior to the incorporation of new habitat (see Supplementary Information).

## 3. Results

A large variation was observed in the magnitude of the impact of the intervention, ranging from quasi extinction (99.4% reduction in occupancy) to a growth in an order of magnitude. Larger interventions lead to impacts of larger magnitudes, both positive and negative; while higher colonization-extinction ratios lead to stronger negative impacts and attenuated positive ones.

We found a series of parameter combinations in the (*θ*, φ) parameter space where the intervention has a negative effect -i.e. reduction in the regional occupancy of the species for which novel habitat is suboptimal (See Figure 2). Under these conditions, adding new habitat erodes species coexistence. This happens at relatively high niche overlap (*θ* >∼ 0.85), indicating interventions have the potential of reducing coexistence within guild members (region above the black line in Figure 2). The size of the parameter space (*θ*, φ) that leads to a reduction in the regional occupancy of one species is reduced with increasing intervention size *H*_*2*_ and increases with growing colonization-extinction ratios. In addition, the minimum *θ* that leads to a negative impact increase with intervention size *H*_*2*_ and is reduced with colonization-extinction ratios. This means that larger interventions favor species coexistence at higher competition levels. However, if these species have higher colonization over extinction ratios it is more likely that the intervention determines a negative effect in coexistence. On the other hand, for a fixed θ, the φ value at which the impact is negative also increases with intervention size and colonization-extinction ratios. This means that larger interventions with species with higher colonization rates require a higher degree of similarity between original and incorporated habitat to have a positive effect in species performance and coexistence (Figure 2).

**Figure 2.**
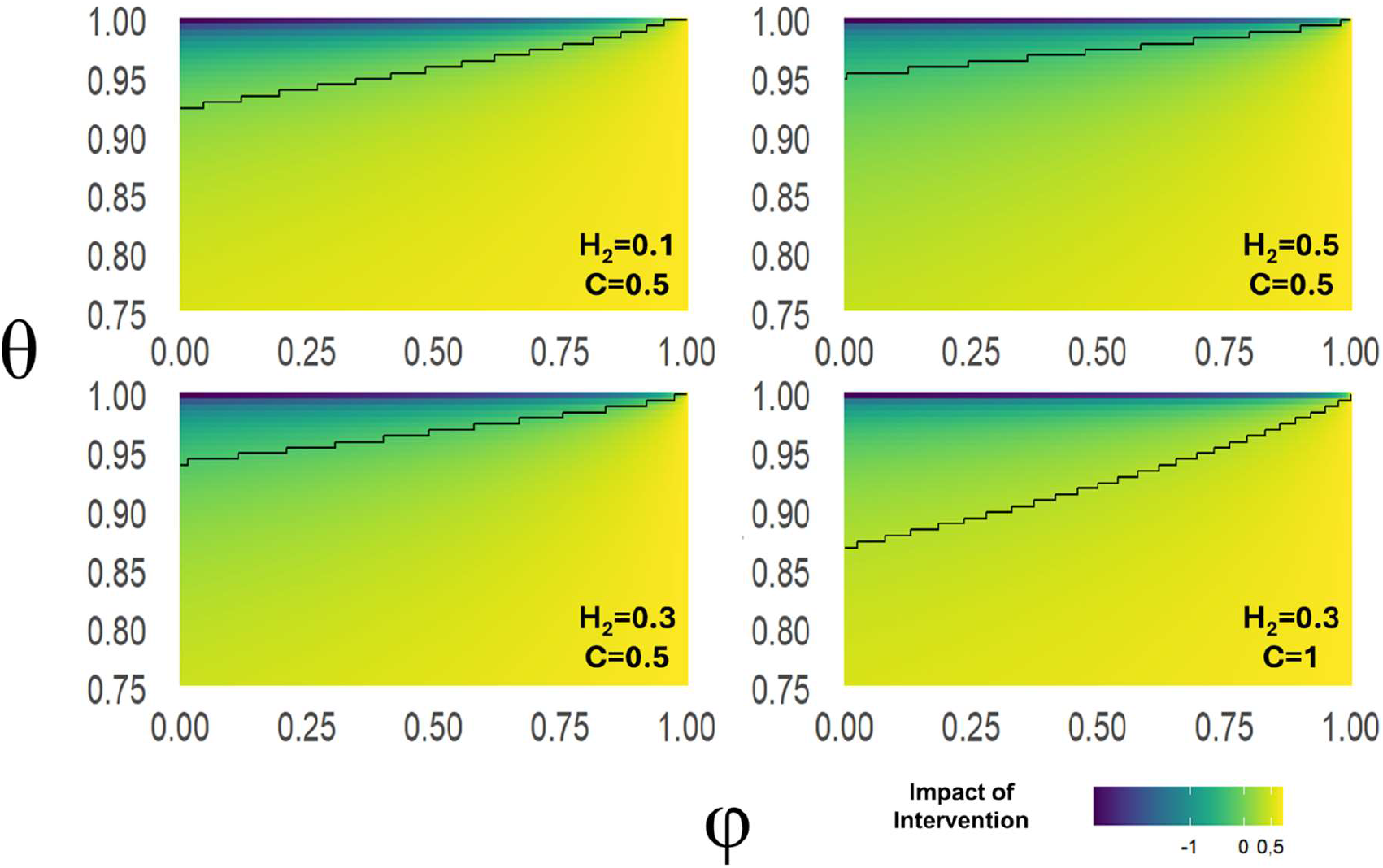
Impact of the intervention (in colors) for different parameter combinations. Horizontal axis represents the fidelity of the intervention (i.e. how much the novel habitat resembles the original one) while the vertical axis represents competition between species. Each panel shows a different combination of added habitat and colonization rates. Black line separates negative impacts (above the line) from positive ones (below the line). Extinction rates are fixed.

## 4. Discussion

Restoration constitutes one of the most relevant tools that human societies have to tackle global change consequences and preserve biodiversity and ecosystem function (Abrams 2018; Dobson 1997, United Nations Decade on Ecosystem Restoration Resolution, 2019). In this analysis, we highlight that suboptimal restoration can foster competitive exclusion of coexisting species, representing a potential double-edged sword for conservation. This shows that a better understanding of the mechanisms connecting restoration interventions with their effects on biodiversity is still required (Cooke et al.,2020; Durant et al., 2019; Kadykalo et al., 2021). In this study we related the addition of novel habitat with the stabilizing and equalizing mechanisms that determine species coexistence (Chesson 2000, 2020). Notably, we identified mechanisms through which a suboptimal intervention, while increasing all species performances, can unbalance the relative performance of species in the landscape, increasing one of the species competitive ability and affecting coexistence at the landscape level (Chesson 2020). Furthermore, we identify the conditions that promote this effect, which are related to groups of species with high niche overlap, a relatively small scale of intervention, and the creation of novel habitats that generate a disproportionate fitness advantage of one group of species.

Our model focuses on the competitive interaction of two species. However, our mean field approach can be conceptually extended to include two sets of competing pools of *n* species (i.e. metacommunity). In this case, the expected species richness of each pool in the metacommunity is the total species richness of the pool times the fraction of occupied patches by the pool (Hanski, 2009). Thus, a diminished occupancy of species pool one results in a species richness reduction at the metacommunity level. Likewise, within a metacommunity there could exist a given set of pairs of competing species to which our model conclusions could be applied, resulting in a generalized diversity loss. When extending our conclusions to a metacommunity context, it is relevant to keep in mind that other important metacommunity processes, such as regional dispersal or high order interactions (Leibold & Chase 2018; Gibbs et al., 2022), could magnify or diminish the negative effects of suboptimal restoration.

Our results show that suboptimal restoration has the potential of negatively impacting species with a relatively high niche overlap, for example ecologically similar species belonging to the same guild (Godoy & Levine, 2014, Bimler et al., 2018). In this sense, species that coexist in neutral or quasi neutral dynamics (with a fitness ratio and niche overlap of 1, right side of Fig. 1) are particularly vulnerable, as inducing small differences in fitness may lead to competitive exclusion. It is expectable that within a metacommunity these interactions of species with complete niche overlap (i.e. neutrality) or nearly complete overlap (i.e. quasi neutrality) exist (Bimler et al., 2018; Souza et al., 2016; Adler et al., 2007; Leibold & McPeek, 2006; Hubbell, 2001). In this sense, restoration strategies that focus on returning the system to a desired, pre-degradation state should be particularly careful in designing interventions that avoid large disruptions in the fitness differences of species. The inclusion of heterogeneity in the attributes of restored patches could avoid this, as the fitness gains could be averaged along the patches.

On the other hand, our results could also be used to think of suboptimal restoration as a useful management tool. The effect that suboptimal restoration can have on coexistence can be used to favor species negatively impacted by human activity or invasive species (Montràs-Janer et al., 2024). By identifying conditions that favor the desired species more than the undesired ones, managers can shift the coexistence regime at the metacommunity scale by generating relatively small interventions that have this particular objective in mind (Hallet et al., 2023). This strategy differs from the attempts to to get the system to a pre-degradation state through the creation of habitat that tries to mimic the original one. An example of this could be the design of restoration interventions which explicitly promote competitive exclusion of invasive species; or interventions that make sure the restored habitat favors those species with declining populations more than it favors their competitors. The challenges associated with using this suboptimal restoration as a tool are associated with identifying which species need to be favored and unfavored, what is their niche overlap, and what habitat attributes will lead to the desired outcomes (Mathakuta et al., 2019; Chinchorro et al., 2019; Aoyama et al., 2022). However, having advanced on this information the addition and preservation of habitats in which endangered species perform better than its competitors is here mechanistically supported as a relevant management strategy.

The advantage held by one of the species in the restored habitat is extended to the original ones by means of mass effect (Brown & Kodric-Brown 1977; Leibold et al., 2004; Leibold & Chase, 2018). Higher abundances of the favored species in the restored habitat lead, via colonization, to higher occupancies in the original habitat. In this sense, it should be noted that interventions aimed at restoring landscape connectivity, for example through the restoration of biological corridors or stepping stones (Gilbert et al, 1998; Crooks & Sanjay 2006; Breckheimer et al., 2014; Ramiadantsoa et al., 2015), could result in intensified positive or negative outcomes.

The transient dynamic from the intervention to the final outcome could take decades to happen. Additionally, if there exists a mismatch between the timescales of competitive exclusion and the colonization of new habitat, a positive trend in diversity could be observed before diversity loss (Aoyama et al. 2022). A more thorough study on the possible dynamics of this transient and its response to variations in the studied parameters could lead to a better understanding of the phenomena. For instance, future studies on these transients could be useful to detect early warning signs that interventions are not having the desired results. Further investigations using stochastic, agent based, or more complex models could be useful in order to get a better grasp of the complexities involved in these scenarios (e.g. Zhang et al., 2024). However, it has to be highlighted that the generality of the modern coexistence theory suggests that our results should be robust to variation in the modeling framework, as long as the considered scenario is preserved (Chesson 2020).

While theoretical ecology has had a meaningful impact in the study of the effects of habitat destruction, it has made fewer contributions to restoration ecology (Perring et al., 2015; Torok & Helm, 2017; Gawecka & Bascompte, 2021; Hallet et al., 2023). Theoretical research is important to understand mechanisms behind observed patterns, generate novel predictions and comprehend counterintuitive results (Marquet et al., 2014; Rossberg et al., 2019). The results of this work are relevant to understand and prevent potentially undesired outcomes of restoration implementations. We identify potential interventions that result in biodiversity loss depending on a small set of parameters, namely the magnitude of niche overlap between present species (for example functional redundancy among species), the fidelity of the intervention (i.e. if and how much the novel habitat is more suitable for some species than others), the colonization-extinction ratios of the species (i.e. species dispersal ranges and regional connectivity), and the amount of habitat restored (i.e. number and size of restored patches). The underlying mechanism behind this observation can be traced back to modern coexistence theory and the balance between the stabilizing and equalizing mechanisms that allow coexistence.

## Acknowledgements

This study was designed and performed in the framework of the European Commission, PONDERFUL Horizon 2020 project (H2020-LC-CLA-2019-2). F.M.U. acknowledges the support and funding of Agencia Nacional de Investigación e Innovación POS_NAC_2021_1_170399. F.M.U. would like to thank David Alonso for relevant insights and feedback. D.C.M was supported by the European Union’s Horizon 2020 research and innovation programme under the Marie Skłodowska-Curie grant agreement No 101062388

## References

Abrams, R. W. (2018). Habitat restoration: A 21st century pursuit. In Reference Module in Earth Systems and Environmental Sciences. Elsevier. 10.1016/b978-0-12-409548-9.10867-x

Adler, P. B., HilleRisLambers, J., & Levine, J. M. (2007). A niche for neutrality. Ecology Letters, 10(2), 95–104. 10.1111/j.1461-0248.2006.00996.x

Albert, J. S., Destouni, G., Duke-Sylvester, S. M., Magurran, A. E., Oberdorf, T., Reis, R. E., Winemiller, K. O., & Ripple, W. J. (2020). Scientists’ warning to humanity on the freshwater biodiversity crisis. Ambio, 50(1), 85–94. 10.1007/s13280-020-01318-8

Aoyama, L., Shoemaker, L.G., Gilbert, B., Collinge, S.K., Faist, A.M., Shackelford, N., Temperton, V.M., Barabás, G., Larios, L. & Ladouceur, E. (2022) Application of modern coexistence theory to rare plant restoration provides early indication of restoration trajectories. Ecological Applications, 32, e2649.

Barabás, G., D’Andrea, R., & Stump, S. M. (2018). Chesson’s coexistence theory. Ecological Monographs, 88(3), 277–303. 10.1002/ecm.1302

Bascompte, J., & Solé, R. V. (1998). Effects of habitat destruction in a prey–predator metapopulation model. Journal of Theoretical Biology, 195(3), 383–393. 10.1006/jtbi.1998.0803

Battisti, C. (2003). Habitat fragmentation, fauna and ecological network planning: Toward a theoretical conceptual framework. Italian Journal of Zoology, 70(3), 241–247. 10.1080/11250000309356524

Bimler, M. D., Stouffer, D. B., Lai, H. R., & Mayfield, M. M. (2018). Accurate predictions of coexistence in natural systems require the inclusion of facilitative interactions and environmental dependency. Journal of Ecology, 106(5), 1839–1852. 10.1111/1365-2745.13030

Breckheimer, I., Haddad, N. M., Morris, W. F., Trainor, A. M., Fields, W. R., Jobe, R. T., Hudgens, B. R., Moody, A., & Walters, J. R. (2014). Defining and evaluating the umbrella species concept for conserving and restoring landscape connectivity. Conservation Biology, 28(6), 1584–1593. 10.1111/cobi.12362

Brown, J. H., & Kodric-Brown, A. (1977). Turnover rates in insular biogeography: Effect of immigration on extinction. Ecology, 58(2), 445–449. 10.2307/1935620

Cabrera, S., Compte, J., Gascón, S., Boix, D., Cunillera-Montcusí, D., Barrero, L., & Quintana, X. D. (2019). How do zooplankton respond to coastal wetland restoration? the case of newly created salt marsh lagoons in La Pletera (NE Catalonia). Limnetica, 38(2), 721–741. 10.23818/limn.38.42

Chesson, P. (2000a). General theory of competitive coexistence in spatially-varying environments. Theoretical Population Biology, 58(3), 211–237. 10.1006/tpbi.2000.1486

Chesson, P. (2000b). Mechanisms of maintenance of species diversity. Annual Review of Ecology and Systematics, 31(1), 343–366. 10.1146/annurev.ecolsys.31.1.343

Chesson, P. (2003). Quantifying and testing coexistence mechanisms arising from recruitment fluctuations. Theoretical Population Biology, 64(3), 345–357. 10.1016/s0040-5809(03)00095-9

Chesson, P. (2018). Updates on mechanisms of maintenance of species diversity. Journal of Ecology, 106(5), 1773–1794. 10.1111/1365-2745.13035

Chesson, P. (2020). Species coexistence. In Theoretical Ecology (pp. 5–27). Oxford University Press. 10.1093/oso/9780198824282.003.0002

Chichorro, F., Juslén, A., & Cardoso, P. (2019). A review of the relation between species traits and extinction risk. Biological Conservation, 237, 220–229. 10.1016/j.biocon.2019.07.001

Cooke, S. J., Rytwinski, T., Taylor, J. J., Nyboer, E. A., Nguyen, V. M., Bennett, J. R., Young, N., Aitken, S., Auld, G., Lane, J.-F., Prior, K. A., Smokorowski, K. E., Smith, P. A., Jacob, A. L., Browne, D. R., Blais, J. M., Kerr, J. T., Ormeci, B., Alexander, S. M., … Smol, J. P. (2020). On “success” in applied environmental research — What is it, how can it be achieved, and how does one know when it has been achieved? Environmental Reviews, 28(4), 357–372. 10.1139/er-2020-0045

Crooks, K. R., & Sanjayan, M. (2006). Connectivity conservation. Cambridge University Press.

Crouzeilles, R., Curran, M., Ferreira, M. S., Lindenmayer, D. B., Grelle, C. E. V., & Rey Benayas, J. M. (2016). A global meta-analysis on the ecological drivers of forest restoration success. Nature Communications, 7(1). 10.1038/ncomms11666

Cuenca-Cambronero, M., Blicharska, M., Perrin, J.-A., Davidson, T.A., Oertli, B., Lago, M., Beklioglu, M., Meerhoff, M., Arim, M. & Teixeira, J. (2023) Challenges and opportunities in the use of ponds and pondscapes as Nature-based Solutions. Hydrobiologia, 1–15.

Dobson, A. P., Bradshaw, A. D., & Baker, A. J. M. (1997). Hopes for the future: Restoration ecology and conservation biology. Science, 277(5325), 515–522. 10.1126/science.277.5325.515

Durant, S. M., Groom, R., Kuloba, B., Samna, A., Muzuma, U., Gadimang, P., Mandisodza-Chikerema, R., Ipavec, A., Mitchell, N., Ikanda, D., & Msuha, M. (2019). Bridging the divide between scientists and decision-makers: How behavioural ecologists can increase the conservation impact of their research? Philosophical Transactions of the Royal Society B: Biological Sciences, 374(1781), 20190011. 10.1098/rstb.2019.0011 Ecosystems and human well-being: Synthesis. (2005).

Fortuna, M. A., Krishna, A., & Bascompte, J. (2012). Habitat loss and the disassembly of mutalistic networks. Oikos, 122(6), 938–942. 10.1111/j.1600-0706.2012.00042.x

Gawecka, K. A., & Bascompte, J. (2021). Habitat restoration in spatially explicit metacommunity models. Journal of Animal Ecology, 90(5), 1239–1251. 10.1111/1365-2656.13450

Gibbs, T., Levin, S. A., & Levine, J. M. (2022). Coexistence in diverse communities with higher-order interactions. Proceedings of the National Academy of Sciences, 119(43). 10.1073/pnas.2205063119

Gilbert, F., Gonzalez, A., & Evans-Freke, I. (1998). Corridors maintain species richness in the fragmented landscapes of a microecosystem. Proceedings of the Royal Society of London. Series B: Biological Sciences, 265(1396), 577–582. 10.1098/rspb.1998.0333

Godet, L., & Devictor, V. (2018). What conservation does. Trends in Ecology & Evolution, 33(10), 720–730. 10.1016/j.tree.2018.07.004

Godoy, O., & Levine, J. M. (2014). Phenology effects on invasion success: Insights from coupling field experiments to coexistence theory. Ecology, 95(3), 726–736. 10.1890/13-1157.1

Gyllenberg, M., & Hanski, I. (1997). Habitat deterioration, habitat destruction, and metapopulation persistence in a heterogenous landscape. Theoretical Population Biology, 52(3), 198–215. 10.1006/tpbi.1997.1333

Hallett, L.M., Aoyama, L., Barabás, G., Gilbert, B., Larios, L., Shackelford, N., Werner, C.M., Godoy, O., Ladouceur, E.R. & Lucero, J.E. (2023) Restoration ecology through the lens of coexistence theory. Trends In Ecology & Evolution.

Hanski, I. (1994). Patch-occupancy dynamics in fragmented landscapes. Trends in Ecology & Evolution, 9(4), 131–135. 10.1016/0169-5347(94)90177-5

Hanski, I. (2009). The theories of island biogeography and metapopulation dynamics. In The Theory of Island Biogeography Revisited (pp. 186–213). Princeton University Press. 10.1515/9781400831920.186

Hanski, I., Moilanen, A., Pakkala, T., & Kuussaari, M. (1996). The quantitative incidence function model and persistence of an endangered butterfly metapopulation. Conservation Biology, 10(2), 578–590. 10.1046/j.1523-1739.1996.10020578.x

Hanski, I., & Ovaskainen, O. (2000). The metapopulation capacity of a fragmented landscape. Nature, 404(6779), 755–758. 10.1038/35008063

Hanski, I., & Thomas, C. D. (1994). Metapopulation dynamics and conservation: A spatially explicit model applied to butterflies. Biological Conservation, 68(2), 167–180. 10.1016/0006-3207(94)90348-4

Hernández-Carrasco, D., Cunillera-Montcusí, D., Antón-Pardo, M., Cañedo-Argüelles, M., Bas-Silvestre, M., Compte, J., Gascón, S., Quintana, X. D., & Boix, D. (2023). Ecological restoration promotes zooplankton network complexity in Mediterranean coastal lagoons. Restoration Ecology, 31(5). 10.1111/rec.13920

Hubbell, S. P. (2011). The unified neutral theory of biodiversity and biogeography (MPB-32). Princeton University Press.

Kadykalo, A. N., Buxton, R. T., Morrison, P., Anderson, C. M., Bickerton, H., Francis, C. M., Smith, A. C., & Fahrig, L. (2021). Bridging research and practice in conservation. Conservation Biology, 35(6), 1725–1737. 10.1111/cobi.13732

Klausmeier, C. A. (2001). Habitat destruction and extinction in competitive and mutualistic metacommunities. Ecology Letters, 4(1), 57–63. 10.1046/j.1461-0248.2001.00195.x

Laurance, W. F., Koster, H., Grooten, M., Anderson, A. B., Zuidema, P. A., Zwick, S., Zagt, R. J., Lynam, A. J., Linkie, M., & Anten, N. P. R. (2012). Making conservation research more relevant for conservation practitioners. Biological Conservation, 153, 164–168. 10.1016/j.biocon.2012.05.012

Leibold, M. A., & Chase, J. M. (2018). Metacommunity ecology, volume 59. Princeton University Press.

Leibold, M. A., Holyoak, M., Mouquet, N., Amarasekare, P., Chase, J. M., Hoopes, M. F., Holt, R. D., Shurin, J. B., Law, R., Tilman, D., Loreau, M., & Gonzalez, A. (2004). The metacommunity concept: A framework for multi-scale community ecology. Ecology Letters, 7(7), 601–613. 10.1111/j.1461-0248.2004.00608.x

Leibold, M. A., & McPeek, M. A. (2006). COEXISTENCE OF THE NICHE AND NEUTRAL PERSPECTIVES IN COMMUNITY ECOLOGY. Ecology, 87(6), 1399–1410. 10.1890/0012-9658(2006)87[1399:cotnan]2.0.co;2

Levins, R. (1969). Some demographic and genetic consequences of environmental heterogeneity for biological control. Bulletin of the Entomological Society of America, 15(3), 237–240. 10.1093/besa/15.3.237

Marquet, P. A., Allen, A. P., Brown, J. H., Dunne, J. A., Enquist, B. J., Gillooly, J. F., Gowaty, P. A., Green, J. L., Harte, J., Hubbell, S. P., O’Dwyer, J., Okie, J. G., Ostling, A., Ritchie, M., Storch, D., & West, G. B. (2014). On theory in ecology. BioScience, 64(8), 701–710. 10.1093/biosci/biu098

Mathakutha, R., Steyn, C., le Roux, P. C., Blom, I. J., Chown, S. L., Daru, B. H., Ripley, B. S., Louw, A., & Greve, M. (2019). Invasive species differ in key functional traits from native and non-invasive alien plant species. Journal of Vegetation Science, 30(5), 994–1006. 10.1111/jvs.12772

Melián, C. J., & Bascompte, J. (2002). Food web structure and habitat loss. Ecology Letters, 5(1), 37–46. 10.1046/j.1461-0248.2002.00280.x

Montràs-Janer, T., Suggitt, A.J., Fox, R. et al. Anthropogenic climate and land-use change drive short- and long-term biodiversity shifts across taxa. Nat Ecol Evol 8, 739–751 (2024). 10.1038/s41559-024-02326-7

Moilanen, A., & Hanski, I. (1995). Habitat destruction and coexistence of competitors in a spatially realistic metapopulation model. The Journal of Animal Ecology, 64(1), 141. 10.2307/5836

Moor, H., Bergamini, A., Vorburger, C., Holderegger, R., Bühler, C., Egger, S., & Schmidt, B. R. (2022). Bending the curve: Simple but massive conservation action leads to landscape-scale recovery of amphibians. Proceedings of the National Academy of Sciences, 119(42). 10.1073/pnas.2123070119

Nations, F. and A. O. of the U. (2019). The State of the World’s Biodiversity for Food and Agriculture: FAO COMMISSION ON GENETIC RESOURCES FOR FOOD AND AGRICULTURE ASSESSMENTS • 2019. Food & Agriculture Org.

Nee, S., & May, R. M. (1992). Dynamics of metapopulations: Habitat destruction and competitive coexistence. The Journal of Animal Ecology, 61(1), 37. 10.2307/5506

Perring, M. P., Standish, R. J., Price, J. N., Craig, M. D., Erickson, T. E., Ruthrof, K. X., Whiteley, A. S., Valentine, L. E., & Hobbs, R. J. (2015). Advances in restoration ecology: Rising to the challenges of the coming decades. Ecosphere, 6(8), 1–25. 10.1890/es15-00121.1

Ramiadantsoa, T., Ovaskainen, O., Rybicki, J., & Hanski, I. (2015). Large-Scale habitat corridors for biodiversity conservation: A forest corridor in madagascar. PLOS ONE, 10(7), e0132126. 10.1371/journal.pone.0132126

Rey Benayas, J. M., Newton, A. C., Diaz, A., & Bullock, J. M. (2009). Enhancement of biodiversity and ecosystem services by ecological restoration: A meta-analysis. Science (New York, N.Y.), 325(5944), 1121–1124. 10.1126/science.1172460

Rossberg, A. G., Barabás, G., Possingham, H. P., Pascual, M., Marquet, P. A., Hui, C., Evans, M. R., & Meszéna, G. (2019). Let’s train more theoretical ecologists – here is why. Trends in Ecology & Evolution, 34(9), 759–762. 10.1016/j.tree.2019.06.004

Ruhí, A., Boix, D., Gascón, S., Sala, J., & Quintana, X. D. (2012). Nestedness and successional trajectories of macroinvertebrate assemblages in man-made wetlands. Oecologia, 171(2), 545–556. 10.1007/s00442-012-2440-7

Souza, A. F., Bezerra, A. D., & Longhi, S. J. (2016). Quasi-neutral community assembly: Evidence from niche overlap, phylogenetic, and trait distribution analyses of a subtropical forest in South America. Perspectives in Plant Ecology, Evolution and Systematics, 23, 1–10. 10.1016/j.ppees.2016.09.006

Sutherland, W. J., Dicks, L. V., Petrovan, S. O., & Smith, R. K. (2021). What works in conservation 2021. Open Book Publishers.

Tao, Y., Hastings, A., Lafferty, K. D., Hanski, I., & Ovaskainen, O. (2024). Landscape fragmentation overturns classical metapopulation thinking. Proceedings of the National Academy of Sciences, 121(20). 10.1073/pnas.2303846121

Tickner, D., Opperman, J. J., Abell, R., Acreman, M., Arthington, A. H., Bunn, S. E., Cooke, S. J., Dalton, J., Darwall, W., Edwards, G., Harrison, I., Hughes, K., Jones, T., Leclère, D., Lynch, A. J., Leonard, P., McClain, M. E., Muruven, D., Olden, J. D., … Young, L. (2020). Bending the Curve of global freshwater biodiversity loss: An emergency recovery plan. BioScience, 70(4), 330–342. 10.1093/biosci/biaa002

Tilman, D. (1994). Competition and biodiversity in spatially structured habitats. Ecology, 75(1), 2–16. 10.2307/1939377

Török, P., & Helm, A. (2017). Ecological theory provides strong support for habitat restoration. Biological Conservation, 206, 85–91. 10.1016/j.biocon.2016.12.024

United Nations. (2019). Resolution A/RES/73/284: United Nations Decade on Ecosystem Restoration (2021–2030). Retrieved from https://undocs.org/A/RES/73/284

Young, T. P., Petersen, D. A., & Clary, J. J. (2005). The ecology of restoration: Historical links, emerging issues and unexplored realms. Ecology Letters, 8(6), 662–673. 10.1111/j.1461-0248.2005.00764.x

Zhang, H., Chase, J. M., & Liao, J. (2024). Habitat amount modulates biodiversity responses to fragmentation. Nature Ecology & Evolution. 10.1038/s41559-024-02445-1

