## Supplementary Information for "Similar won’t make it: Coexistence, habitat availability and suboptimal restoration"

To calculate the values of niche overlap, $\rho$, in the context of the model, it is first necessary to determine the values of the intra- and interspecific competition coefficients. These coefficients represent the effects of an individual from the same or different species on the per capita growth rate. We can extend this reasoning assuming that each patch can be occupied only by one individual (see Tilman 1994). Thus, it is necessary to calculate the slopes resulting from plotting the per capita growth rate against the fractions of patches occupied by each set of species, as shown in Figure S1.

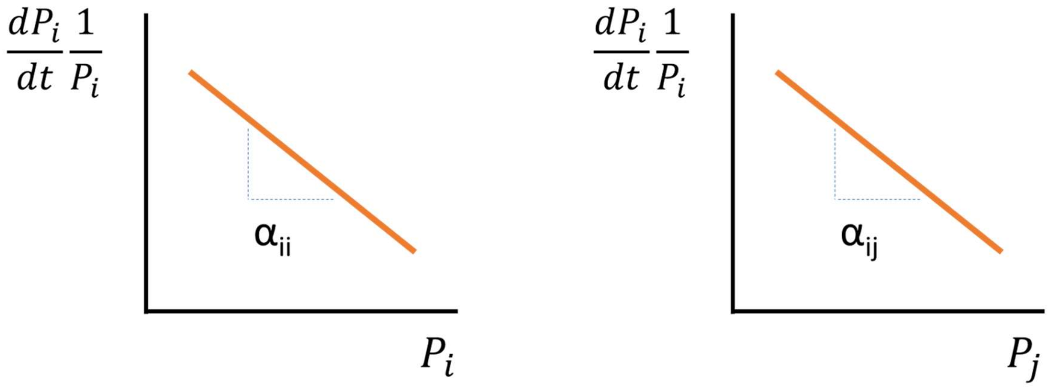


**Figure S1**: Slopes result of plotting per capita growth rates in the fraction of occupied patches against the fraction of occupied patches allows to obtain Lotka-Volterra competition coefficients.

Given that we are trying to find the niche overlap in the original habitat, we can determine $H_1$=1 and $H_2$=0, so $P_{21}$ and $P_{22}$ are also 0.

$$\begin{cases} \dfrac{dP_1}{dt} = cP_1(1 - P_1 - \theta P_2) - eP_1 \\ \\ \dfrac{dP_2}{dt} = cP_2(1 - P_2 - \theta P_1) - eP_2 \end{cases}$$

So:

$$\begin{cases} \dfrac{dP_1}{dt}\dfrac{1}{P_1} = c(1 - P_1 - \theta P_2) - e \\ \\ \dfrac{dP_2}{dt}\dfrac{1}{P_2} = c(1 - P_2 - \theta P_1) - e \end{cases}$$

Reordering the terms:

$$
\begin{cases}
\dfrac{dP_1}{dt}\dfrac{1}{P_1} = -cP_1 + c - c\theta P_2 - e \\
\\
\dfrac{dP_2}{dt}\dfrac{1}{P_2} = -cP_2 + c - c\theta P_1 - e
\end{cases}
$$

Given our model assumes symmetric competition in the original habitat, the terms of intra and interspecific Lotka-Volterra coefficients are the same:

$$\alpha_{12} = \alpha_{21} = -c\theta$$

$$\alpha_{11} = \alpha_{22} = -c$$

And given that (Chesson 2020):

$$\gamma_{ij} = \frac{\alpha_{ij}}{\alpha_{ji}}$$

$$\rho = \sqrt{\gamma_{ij} \cdot \gamma_{ji}}$$

Then:

$$\rho = \theta$$
